# If it’s there, could it be a bear?

**DOI:** 10.1101/2023.01.14.524058

**Authors:** Floe Foxon

**Affiliations:** Folk Zoology Society – PO Box 97014, Pittsburgh, PA, 15229, USA

## Abstract

It has been suggested that the American black bear (*Ursus americanus*) may be responsible for a significant number of purported sightings of an alleged unknown species of hominid in North America. Previous analyses have identified correlation between ‘sasquatch’ or ‘bigfoot’ sightings and black bear populations in the Pacific Northwest using ecological niche models and simple models of expected animal sightings. The present study expands the analysis to the entire US and Canada by regressing sasquatch sightings on bear populations in each state/province while adjusting for human population and forest area in a generalized linear model. Sasquatch sightings were statistically significantly associated with bear populations such that, on the average, every 1, 000 bear increase in the bear population is associated with a 4% (95% CI: 1%–7%) increase in sasquatch sightings. Thus, as black bear populations increase, sasquatch sightings are expected to increase also. On the average, across all states and provinces in 2006, after controlling for human population and forest area, there were approximately 5, 000 bears per sasquatch sighting. Based on statistical considerations, it is likely that many supposed sasquatch are really misidentified known forms. If bigfoot is there, it may be a bear.

## Introduction

The United States and Canada feature nearly 20 million km^2^ of land, hosting hundreds of mammal species in its woodlands, prairies, boreal forests, and along its coasts (Kays & Wilson 2009). Due to this size and diversity, it is unlikely that the North American faunal catalogue is complete, with numerous insects and possibly small mammals remaining to be discovered. While most scientists believe that the existence of a large hominid species in North America, as-yet unrecognised in conventional science, is not likely (King & Greenwell 1983), some authors have entertained this possibility in the field of research called ‘hominology’.

Hominology, as formulated by Boris Porshnev and Dimitri Bayanov, alleges that hairy, bipedal primates not recognised in conventional zoology *do* exist and are relicts of *Homo neanderthalensis*, banished to the wild through competition with *H. sapiens* (Bayanov 2012). One such mystery hominid reported in North America is variously dubbed ‘sasquatch’ after West Coast First Nations tradition (Heuvelmans 1986), ‘bigfoot’ to Westerners, and the *nomen dubium Gigantopithecus canadensis* to cryptozoologists who believe that rather than a hominin, this animal is a relict Pleistocene pongid (Heaney 1990). Purported sasquatch sightings number in the thousands (Bigfoot Field Researchers Organization 2023), making bigfoot a prominent anthrozoological phenomenon.

Hominologists describe six lines of evidence to support the above claims (Bayanov 2012): (1) descriptions of wild men in the natural history texts of ancient Roman and Arab philosophers and medieval Europeans; (2) folklore and mythology; (3) ancient and medieval art; (4) footprints, tracks, etc.; (5) photographs; and (6) eyewitness testimony. These are all categories of indirect evidence (i.e., testimonial and circumstantial).

Concerning (1), while some legendary animals in ancient texts were likely inspired by encounters with real forms (e.g., the Greco-Roman griffin and its probable inspiration in fossil ceratopsians (Mayor 1991)), the existence of other beings catalogued in ancient and medieval bestiaries have not been treated with such conviction. Similarly concerning (2), while various cross-culture traditions, myths, and legends have been identified in association with mystery hominids including sasquatch (Bayanov 2011), many characters in folklore such as magical witches and sorcerers do not correspond to counterparts in reality (Colarusso 1983). In hominology, folklore is interpreted as the emotional expression of lived experiences (Bayanov & Bourtsev 1976), therefore sasquatch traditions are seen as evidence for sasquatch existence. However, many of these folkloric references are extremely heterogeneous in their descriptions of mystery hominids.

Concerning (3), ‘hairy man’ pretroglyphs in the Tule River Tribe Reservation, approximately 1,000 years old, have been interpreted as pictorial representations of bigfoot (Strain 2012). However, other Native American petroglyphs depicting animals such as humanoid frogs have not received such literal interpretations, which may indicate a kind of confirmation bias in hominology research.

Concerning (4), numerous casts and measurements of tracks and footprints attributed to sasquatch have been presented (Napier 1976, ‘Tables’), some apparently featuring primate-like dermal ridge patterns, sweat pores, and sole pads (Krantz 1983; Cachel 1985). These have been criticised as hoaxes constructed with modelling clay and the ‘latex-and-kerosene expansion method’ of preserving details of [human] footprints while greatly increasing their size (Baird 1989; Bodley 1988). It has been suggested that features such as ‘sweat pores’ are casting artefacts such as air bubbles (Freeland & Rowe 1989). Genetic and microscopic analyses of supposed hairs, faeces, and other specimens attributed to sasquatch have been variously identified as synthetic fiber (Winn 1991; Somer 1989), or material from known forms such as cervids, bovines, and ursids (Federal Bureau of Investigation 2019; Bryant & Trevor-Deutsch 1980; Coltman & Davis 2005; Sykes et al. 2014; Hart 2016b,a).

Concerning (5), the ‘Patterson-Gimlin film’ is perhaps the ‘best’ photographic evidence for sasquatch. This notorious 16 mm motion picture purportedly depicts an unknown hominid over six feet tall in California (Munns 2014; Kelsey 2022; Discovery 2022). The film apparently was not spliced or edited (Munns & Meldrum 2013), but many have noted the imposing likelihood that the film subject is a suited actor.

This leaves (6), the testimony of thousands of eyewitnesses. The aim of the present study is to better understand what thousands of people see when they report a sasquatch encounter. Besides hoaxes, the leading hypothesis among skeptics is that when eyewitnesses report seeing bigfoot, they are actually seeing bears (Nickell 2013). To understand how this can be the case, it is necessary in the following to describe the characteristics of bears in North America, and how these are similar to purported characteristics of sasquatch.

North America features three bear species, chiefly among them being the American black bear (*Ursus americanus*). The black bear is a large tetrapod (lengths up to 6 feet or more and masses up to 300 kg), with a pelage of various shades of black, brown, red, blue, blond, and white (Burton 1998). Morphologically, the black bear is stocky with strong musculature around its thick neck, shoulders, and legs (Clark et al. 2021). The tail is short and therefore may be difficult to notice.

The geographic distribution of the black bear is widespread, ranging all across Canada and Alaska, the East and West coasts of the United States, south into Mexico, and with various isolated populations in-between (Beer & Morris 2004). Although the present distribution is much diminished from the historical range, the black bear has experienced a notable expansion since the 1990s (Scheick & Mccown 2014). Black bears may inhabit scrub, swamps, and mountains, but are primarily found in forest and woodland (Beer & Morris 2004; Burton 1998).

Black bears have opportunistic forraging habits; being omnivores, they consume various plant material (including berries, fruits, nuts, and grasses), other mammals (neonate ungulates), fish, invertebrates, and carrion (Beer & Morris 2004). This opportunism means habituated and food-conditioned bears can and will approach people and buildings in search of anthropogenic food sources such as agriculture and garbage (Herrero 2018, Chapter 4).

Although this dietary range adaptation has allowed black bears to remain widespread (Clark et al. 2021), it is also the primary cause of confrontation between humans and bears in areas such as national parks (Herrero 2018, Chapter 4). Black bears are easily food conditioned (Brown 1993, Chapter 4), and because of this they are considered pests in much of North America (Burton 1998).

The presence of the black bear is indicated by such signs as overturned logs and broken twigs and branches (Burton 1998), overturned surface stones, tracks, hairs, and claw and tooth marks left on trees (Herrero 2018, Chapter 4). These bears are solitary and active at night, and may be heard vocalising through grunts, growls, woofs, and howls (Beer & Morris 2004), as well as snorts, roars, coughs, and squeaks, all indicative of emotion (Brown 1993, Chapter 4). Black bears also have a pronounced odor (Brown 1993, Chapter 3). They are inquisitive (Brown 1993, Chapter 4) and intelligent animals (Vonk et al. 2012), capable of forming natural concepts at concrete, intermediate and abstract levels. Bears in general are plantigrade and are known to stand upright and walk on their hind legs; some bear species are known to ambulate over 400 meters bipedally (Brown 1993, Chapter 4). Black bears are also strong swimmers, outstanding climbers, and can run at speeds of up to 13 meters per second (Brown 1993, Chapter 4).

It is precisely the above characteristics of the black bear that make this animal a likely candidate for many purported sasquatch sightings (Nickell 2013). Indeed, Brown (1993, Chapter 5) lists numerous lines of evidence connecting sasquatch traditions with bears, including reported size, strength, and speed; bipedal locomotion; leaving human-or ape-like footprint outlines; tracks found and encounters occurring in bear habitats; and having the same colouration.

Cryptozoologist Ivan Sanderson (1967, Chapter 18) also described sasquatch as “forest dwellers”, sightings of which fall within the distribution of the Earth’s vegetational belts (i.e., where bears and bear foods are found). Other reported characteristics of sasquatch include distinctive smell and various vocalisations; eyes “like a bear’s”; “reddish”, “dark brown”, “light brown”, and other fur colours; consumption of berries and other vegetation; rolling boulders; and raiding of houses for food (Sanderson 1967, Chapters 2–4), are all consistent with black bears in North America, as described above. Of course, an alternative interpretation of these correlates is that the sasquatch is an animal distinct from the black bear but is similar to it in many ways, including habitat and the characteristics described previously.

Existing studies have explored the possible link between the American black bear and bigfoot. Blight (2005) used probabilistic models to examine the relationship between sasquatch sightings and black bear populations in the Pacific Northwest (PNW; including Alaska, Montana, Oregon, Washington, Northern California, and Idaho) and the rest of the US. The method used was a simple calculation of the bivariate correlation coefficient between sasquatch sighting frequency and black bear population density. That study identified positive correlation between black bear populations and sasquatch sightings. One criticism of this study is that the test used assumed a Gaussian distribution for count data. Futhermore, only the US was considered, despite sasquatch sightings being considerably more widespread across the entire US and Canada.

In another study, Lozier et al. (2009) identified a high degree of overlap in predicted geographic distributions for black bears and sasquatch. The method used was to compare results from ecological niche models (ENMs) for each of sasquatch and black bears in the PNW. ENMs are tools used in conservation to predict geographic ranges of animals; they take as input georeferenced data (i.e., events with geographic coordinates) and environmental data layers constructed from bioclimatic variables (e.g. annual mean temperature), and output a predicted geographic distribution for the animal (Sillero et al. 2021). Although the purpose of the study by Lozier et al. (2009) was to highlight potential flaws in these models when errors are introduced by inaccurate specimen identities and georeferencing, the analysis is supportive of a statistical association between sasquatch and black bears; the authors state explicitly that the predicted sasquatch distribution may be seriously biased if many (or all) of the sightings represent misidentified black bears, hence the overlap. The Lozier et al. (2009) analysis considered only data from the PNW (Washington, Oregon and California).

The present study builds upon the works of Blight (2005) and Lozier et al. (2009) by using more recent data, by expanding the analysis to the entire US and Canada, and more appropriately assuming a count data distribution as opposed to a Gaussian distribution. Statistical methods (regression modelling) are used to test the hypothesis that sasquatch sightings and populations of black bears are associated, while controlling for potential confounding by human population and forest area.

## Methods

The number of black bears in each US state and Canadian province were sourced from Table 1. of Spencer et al. (2007). To the author’s knowledge, this represents the latest peer-reviewed, publicly available collection of black bear population data, providing population estimates in each state/province for the year 2006. Because Delaware, Hawaii, Illinois, Indiana, Iowa, Kansas, Nebraska, North Dakota, and South Dakota have no known breeding populations of black bear, the black bear populations were assumed to be zero in these states. Other species of bear were not considered, as their impact on the result is likely to be negligible; black bears are by far the most abundant species, numbering over two times that of all other bear species combined (Garshelis et al. 2021).

**Table 1.** Regression Model Parameters (* *p <* 0.05, ** *p <* 0.0001)

| Variable | Regression Coefficient ± Standard Error |
| --- | --- |
| Intercept | $0.5 \pm 0.2$ * |
| Black Bear Population | $(4 \pm 2) \times 10^{-5}$ * |
| Human Population | $(1.2 \pm 0.2) \times 10^{-7}$ ** |
| Forest Area (km<sup>2</sup>) | $(-7 \pm 3) \times 10^{-6}$ * |

The number of sasquatch sighting reports in each US state and Canadian province were sourced from the Bigfoot Field Researchers Organization’s Geographic Database of Bigfoot/Sasquatch Sightings & Reports (2023). These data consist of eyewitness testimonials, mostly from the second half of the 20^th^ century to the present. To maximise the validity of the analysis, only sightings in the year 2006 were considered such that bear populations and sasquatch sightings were date-matched in these analyses.

Human population statistics for each US state and Canadian province in the year 2006 were obtained from the United States Census Bureau (2021) and Statistics Canada (2022), respectively. Forest area estimates (in km^2^) for each US state in 2006 were sourced from the USDA Forest Service (2006), and estimates for Canadian provinces were sourced from World Atlas (Sawe 2017). For choropleth map plotting, a geojson map of the US and Canada was sourced from Cartograhy Vectors (2022).

Four US states (Rhode Island, Texas, Wisconsin, and Wyoming) and four Canadian provinces (Alberta, Newfoundland and Labrador, Northwest Territories, and Nova Scotia) were missing data on bear population. These were necessarily excluded from analyses.

To test the hypothesis that sasquatch sightings and populations of black bears are associated, a regression model was implemented. This model regressed the number of sasquatch sighting reports in each state/province in 2006 (the dependent/response variable) on the black bear population in each state/province in 2006 (the independent/predictor variable). An intercept term was included to model the possibility of sasqautch sightings in states with no bears (i.e., to model the value of the response variable when the predictor equals zero).

Two more variables were included in the model to reduce confounding and increase the internal validity. These variables were the human population and forest area in each state/province. The more people in a given state/province, the more likely a sighting is to occur. To account for this, the human population in each state/province in 2006 was included as an independent variable in the model. Similarly, the area of forest land in each state/province is likely to impact the number of sasquatch sightings due to the relationship between area and population density, and because black bears are primarily found in forest and woodland (Beer & Morris 2004; Burton 1998).

Consequently, the total amount of forest area (in km^2^) in each state/province was also included as an independent variable in the model.

Because these data (number of sightings) are count data (i.e., countable quantities), it was necessary to use a model appropriate for count data. The model used was a generalized linear model assuming a Negative Binomial distribution and the log link function, which are appropriate for count data. Variations on this model design (e.g., instead assuming Gaussian and Poisson distributions; including interaction terms and random effects; and excluding the intercept term) were investigated in previous exploratory analyses (Foxon 2023b,c). In those exploratory analyses, model fits were assessed by the root mean square error (RMSE; lower is better) and log-likelihood (higher is better). Gaussian models provided the lowest RMSE but it is generally known that Guassian approximations do not correctly describe count data (Lass et al. 2021). Interaction terms were not statistically significant and/or provided poorer model fits to the data, and so were not included in the final model. The inclusion of random intercepts for each state/province overfit the model because only one timepoint is used (i.e., just 2006). The final model in the present study (Negative Binomial, log link, and intercept) provided the highest log-likelihood and is arguably the most appropriate model for these data for the reasons described above.

Indeed, statistical models using negative binomial regression techniques describing the abundance of various animals species have been successfully developed in previous studies in the context of conservation biology, with high predictive power for some species (Pearce & Ferrier 2001; Pradhan & Leung 2006; Acevedo et al. 2014).

All analyses were performed in Python 3.8.16 with the packages Numpy 1.21.5, Pandas 1.5.2, Scipy 1.7.3, Statsmodels 0.13.5, and Plotly 5.9.0. All code and data are available in the online Supplementary Information (Foxon 2023d). Statsmodels uses the iteratively reweighted least squares (IRLS) method for generalized linear models to reduce the impact of outliers.

This work uses only publicly available secondary data on animal subjects. It did not involve primary (*in vivo*) animal research and so is exempt from ARRIVE guidelines.

## Results

Figure 1 shows choropleth maps for the number of sasquatch sightings, black bear populations, human populations, and forest area in the United States and Canada in 2006. At first glance, there are locations such as Florida with very many sasquatch sightings but relatively few bears (at least compared to some other states/provinces). This appears to be inconsistent with the hypothesis that sasquatch sightings and populations of black bears are associated. However, this simple comparison of bears to bigfoot is confounded by the human population and forest area in each state/province. When considering the three maps of bears, people, and forest area simultaneously, the possible associations become more apparent. For example, Florida has relatively few bears and relatively little forest area, but also a relatively large number of people to make sightings. It may be that the combination of fewer bears and less forest area but more people explains the higher number of sightings. Thus, the only way to investigate the possible association between sasquatch sightings and populations of black bears is by considering the three predictors (bears, people, and forest area) simultaneously, which is the purpose of the model.

**Figure 1.**
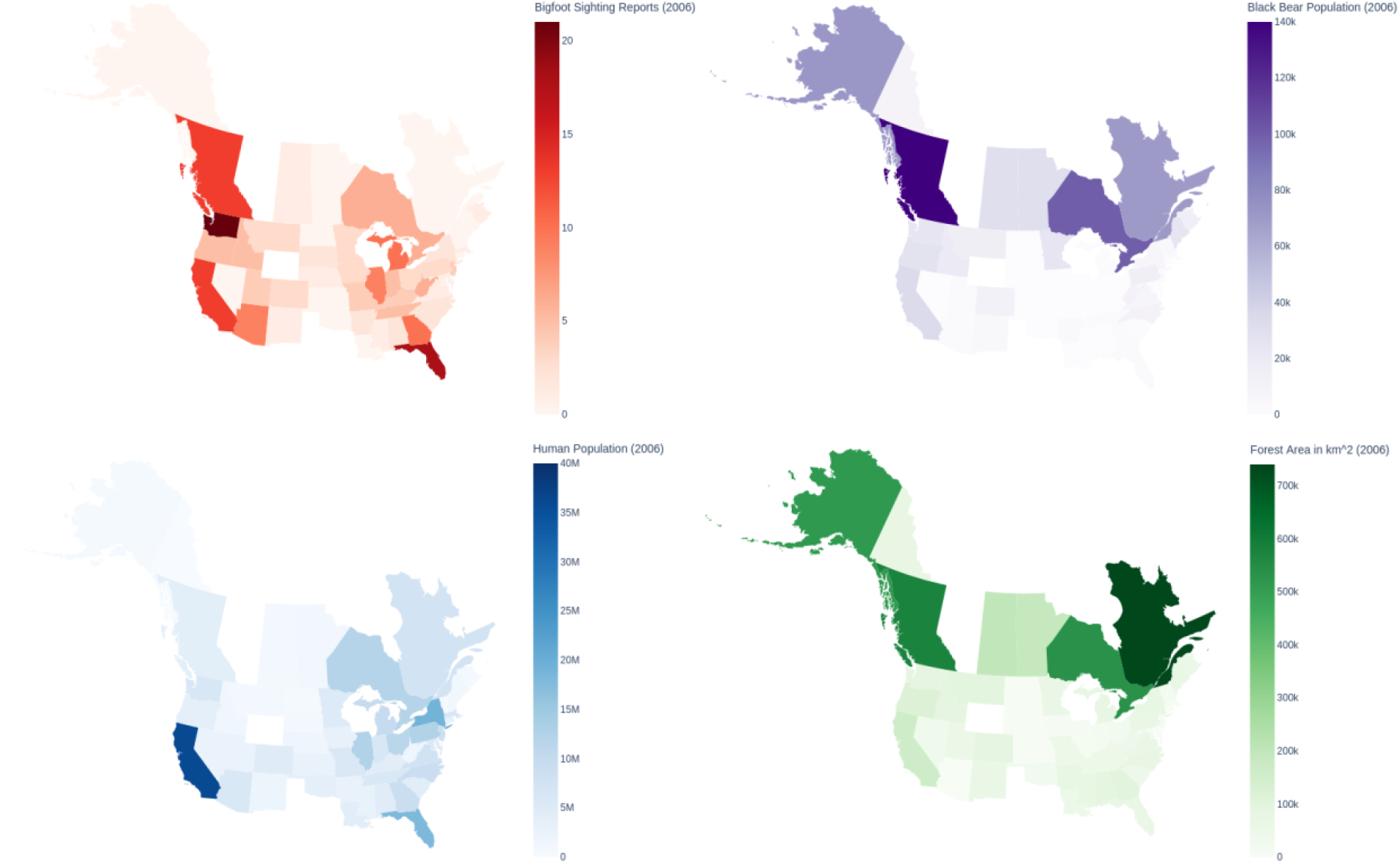
Choropleth maps for sasquatch reports, black bear (*Ursus americanus*) populations, human populations, and forest area across the United States and Canada.

The results of the model regressing bears, people, and forest area on sasquatch sightings in 2006 are described in Table 1. In this model, black bear population was associated with sasquatch sightings such that, on the average, every 1, 000 bear increase in the bear population is associated with a 4% (95% CI: 1%–7%) increase in sasquatch sightings.

Thus, as black bear populations increase, sasquatch sightings are expected to increase also. On the average, across all states and provinces in 2006, after controlling for human population and forest area, there were approximately 5, 000 bears per sasquatch sighting (note that being an average across states/provinces, this estimate is not accurate for each individual state/province). The RMSE of the model (19) was high relative to the number of bigfoot sightings in 2006 in each state/province (range: 0–21).

## Discussion

In the present study, a model was used to investigate the hypothesis that sasquatch sightings are associated with black bear (*Ursus americanus*) populations across the US and Canada. From this model, a positive association was identified between sasquatch sightings and black bears such that, after adjusting for human population and forest area, one sasquatch sighting per year is expected for every 5, 000 bears in North America.

While correlation does not equal causation, perhaps the most parsimonious interpretation of these findings is that many supposed sasquatch sightings in North America may be explained as misidentified black bears. This is logical, because as noted by Nickell (2013) and Brown (1993, Chapter 5), bears and sasquatch share many characteristics in habitat, appearance (e.g., size, hair/fur coverage, and colouration), attributes (e.g., speed and strength), and behaviour (e.g., bipedal locomotion and curiosity). Nickel also notes that poor viewing conditions, non-expert observation, and expectant attention (i.e. ‘seeing’ what one anticipates) could explain why some people might confuse bears for mystery hominids.

Of course, hominologists may point to the possibility that sasquatch are animals distinct from bears but which live in the same places and look and act similarly. However, in the absence of autoptical evidence (i.e., not testimonial or circumstantial), there is not at present strong scientific support for the existence of sasquatch as a mystery hominid.

Although preliminary, these findings suggest that sasquatch sightings may have some utility in bear conservation efforts as a form of citizen science. Data from amateurs/non-professionals are already in use in conservation biology/ecology with great success while also being low in cost (Sumner et al. 2019; MacPhail & Colla 2020; McKinley et al. 2017). It may be that collaboration between bear conservationists and bigfoot field researchers could lead to more recent and accurate estimates of bear population and geographic distribution across North America, improving occupancy estimation by reducing non-detection error (false negatives).

Indeed, cryptozoological anecdotes, in plurality, may well contain legitimate zoological insights (Opit 2017). Paxton (2009) has found that in the context of eyewitness reports of unidentified, large marine animals, cryptozoological anecdotes are amenable to scientific analysis and can be considered data. However, caution must be taken; false-positive (species misidentification) observations can cause overestimation of the distribution or abundance of a species (Costa et al. 2015), as in the case of the Eurasian lynx in the Alps (Molinari-Jobin et al. 2012), the white marlin in the western North Atlantic (Beerkircher et al. 2009), and perching birds in South America (Gorleri & Areta 2022; Gorleri et al. 2023). The solution may be to utilise more advanced models that simultaneously account for both false negative and false positive identification (Miller et al. 2011).

The primary limitation of this study is the potential for residual confounding by factors other than bear population, human population, and forest area. For example, people living, hunting, or hiking in forest areas may also be misidentified as sasquatch. Furthermore, it is important to note that sasquatch sightings have been reported in states with no known breeding black bear populations. Indeed, the intercept term of the model was positive, implying that there are sasquatch sightings even when the number of bears equals zero. Although this may be interpreted as evidence for the possible existence of an unknown hominid in North America, it may also explained by misidentifiation of other animals (including humans), among other possibilities. Still, the present study does not prove that all sasquatch sightings are bear sightings (nor was this study designed for that purpose). Due to these and other limitations specified in the data sources, the findings of the present study must be interpreted as only approximate and not exact (Indeed, as noted above the RMSE was high). Further research and more data are necessary to improve these estimates.

Strengths of this study include the use of a quantitative model that controlled for possible confounding by human population size and forest area in each US state and Canadian province. Furthermore, positive associations between bear populations and bigfoot sightings were also identified in previous exploratory analyses with variations on the model design and data (Foxon 2023b,c), which may suggest that these findings are robust. These findings are also in agreement with the results of previous studies by Blight (2005) and Lozier et al. (2009), which similarly indentified positive associations between bears and bigfoot using different methods. Therefore, the association is likely to be real. Blight (2005) reported only weak positive correlation between bears and bigfoot, whereas the association in the present study is rather more strong. This may be because Blight (2005) assumed a continuous probability distribution (as opposed to a count distribution), did not consider Canada, did not date-match the data, and did not control for forest area. The findings of the present study were closer to those of Lozier et al. (2009), who reported a high ecological niche model overlap statistic (i.e., a strong association between bear and sasquatch geographic distributions). Another stength is that in the present study, the analyses considered the entire US and Canada, whereas the previously-published works only considered the US or Pacific Northwest. Furthermore, the data in the present study were date-matched (in 2006) to provide more accurate estimates of the association.

In conclusion, if bigfoot is there, it may be a bear.

## Acknowledgements

The author thanks Julie Sheldon and Rahul Raveendran for helpful comments and suggestions.

## Data, script, code, and supplementary information availability

Data and code are available online: https://doi.org/10.17605/OSF.IO/AV3G2

